# A Novel Transfer Learning Approach for Toxoplasma Gondii Microscopic Image Recognition by Fuzzy Cycle Generative Adversarial Network

**DOI:** 10.1101/567891

**Authors:** Sen Li, Aijia Li, Diego Alejandro Molina Lara, Jorge Enrique Gómez Marín, Mario Juhas, Yang Zhang

## Abstract

*Toxoplasma gondii*, one of the world’s most common parasites, can infect all types of warm-blooded animals, including one-third of the world’s human population. Most current routine diagnostic methods are costly, time-consuming, and labor-intensive. Although *T.gondii* can be directly observed under the microscope in tissue or spinal fluid samples, this form of identification is difficult and requires well trained professionals. Nevertheless, the traditional identification of parasites under the microscope is still performed by a large number of laboratories. Novel, efficient and reliable methods of *T.gondii* identification are therefore needed, particularly in developing countries. To this end, we developed a novel transfer learning based microscopic image recognition method for *T.gondii* identification. This approach employs Fuzzy Cycle Generative Adversarial Network (FCGAN) with transfer learning utilizing knowledge gained by the parasitologists that Toxoplasma is in banana or crescent shaped form. Our approach aims to build connection between micro and macro associated objects by embedding fuzzy C-means cluster algorithm into Cycle Generative Adversarial Network (Cycle GAN). Our approach achieves 93.1% and 94.0% detection accuracy for 400X and 1000X Toxoplasma microscopic images respectively. We show the high accuracy and effectiveness of our approach in the newly collected unlabeled Toxoplasma microscopic images, comparing to other current available deep learning methods. This novel method for Toxoplasma microscopic image recognition will open a new window for developing cost-effective and scalable deep learning based diagnostic solution, potentially enabling broader clinical access in developing countries.

## 1 Introduction

*Toxoplasma gondii* is a ubiquitous, single cell protozoan parasite that can infect all warm-blooded animals as well as one-third of human population worldwide [Khan and Grigg, 2017]. Most infections in humans are life-long and several studies have suggested that such infections may contribute to severe neurological and psychiatric symptoms. That makes diseases caused by *T.gondii* is one of the biggest health care problems globally. Diagnosis of *T.gondii* is typically performed by testing blood or other body fluids for antibodies or the parasite’s DNA [Burrells *et al.*, 2018]. Large number of laboratories still perform identification of *T.gondii* in tissue or spinal fluid samples under the microscope. However, the microscopic detection and quantification of *T.gondii* is time-consuming, labor-intensive and needs well trained professionals. Moreover, the intensity of illumination, image brightness, contrast level and background of staining, have large variations in different Field of View (FoV), shown in Figure 1. In addition, *T.gondii* is found both inside and outside of nucleated cells with separated and aggregated form, which makes *T.gondii* recognition task more challenging.

**Figure 1:**
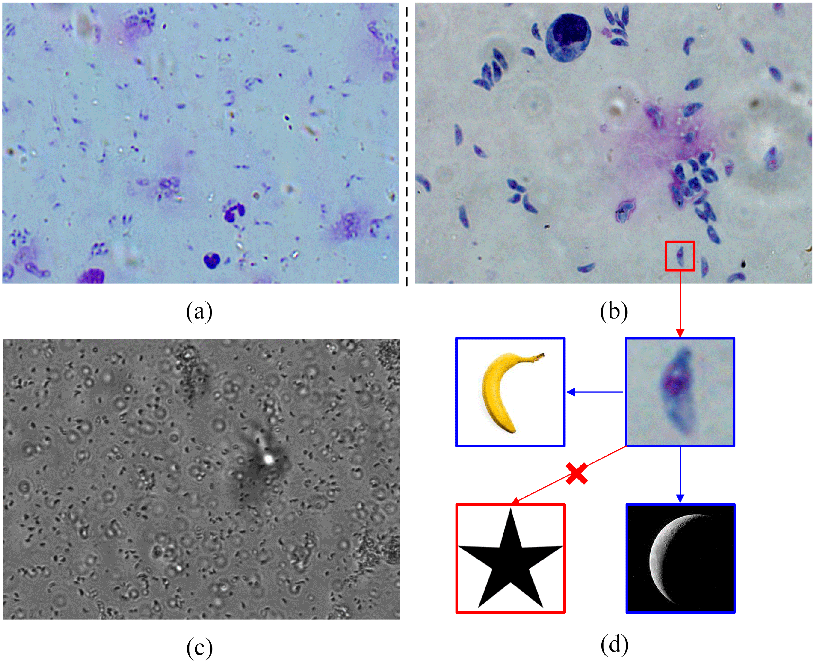
Typical variations of color, illumination, background in different fields of view and microscope magnification. (a) Stained *T.gondii* images from different slides captured by a microscope with 400X magnification; (b) Stained *T.gondii* images from different slides captured by a microscope with 1000X magnification; (c) Unstained *T.gondii* image captured by a microscope with 100X magnification; (d) Training image data with similar macro objects source (banana and crescent) and different macro object (star).

Recently, deep learning technology has significantly improved the efficiency and accuracy in macroscopic computer vision task, thereby attracting considerable attention in microscopic image analysis [Christiansen *et al.*, 2018; Andrea *et al.*, 2018; Baris *et al.*,; Xing *et al.*, 2017]. Sivaramakrishnan [Sivaramakrishnan *et al.*, 2018] evaluated the performance of pre-trained Convolutional Neural Network (CNN) as feature extractor in the classification of parasites and host cells, which improved the infection disease screening. Furthermore, Mehanian [Mehanian *et al.*, 2017] developed a computer vision system with deep learning to identify malaria parasites under the microscope in the field-prepared thick blood films. However, most of the existing deep learning methods on parasite analysis are under supervised learning framework, which requires many well-trained professionals to label a number of image datasets. Furthermore, labeling, annotating, and sorting the output data is time consuming, costly, and labor-intensive. This severely limits their scalability in practical applications.

Moreover, current existing parasite recognition models are limited mainly to malaria. Here, we propose a deep learning method for *T.gondii* recognition. To improve learning efficiency in the task, we employed transfer learning strategy by leveraging knowledge from the parasitologists that *T.gondii* is in crescent shaped form, which is similar to banana. Even more, the shape of aggregated parasites resembles the images of a bunch of bananas, where host cells are significantly different. It is assumed that microscopic object has an inherent connection with macroscopic world. Based on this assumption, we designed a Micro-Macro Associated Object Pulling (M2AOP) strategy and propose a Fuzzy Cycle Generative Adversarial Network (FCGAN) for *T.gondii* recognition. This strategy embeds fuzzy C-means cluster algorithm [Bezdek, 1973] into Cycle Generative Adversarial Network (Cycle GAN)[Zhu *et al.*, 2017], which can learn degrees of membership belonging to each cluster (class) point. Then the degrees are utilized as the translated coefficient to replace the microscopic and macroscopic domain labels in Cycle GAN when the microscopic images are inductive transferred into macroscopic world. FCGAN can exploit more discriminative information using a third associated object to asymmetrically pull *T.gondii* and host cell samples.

We tested two *T.gondii* image datasets consisting of totally 13,138 microscopic images with magnification of 400X, and 14,992 microscopic images with magnification of 1000X. A number of experiments were conducted demonstrating the effectiveness of our FCGAN method with high accuracy and precision.

## 2 Material and Methods

Our proposed method is empowered by a Micro-Macro Associated Object Pulling strategy for microscopic parasite recognition. We utilized Fuzzy Cycle Generative Adversarial Network (FCGAN), shown in Figure 2, where cycle-consistency and fuzzy discriminator loss are optimized along with a fuzzy C-means algorithm.

**Figure 2:**
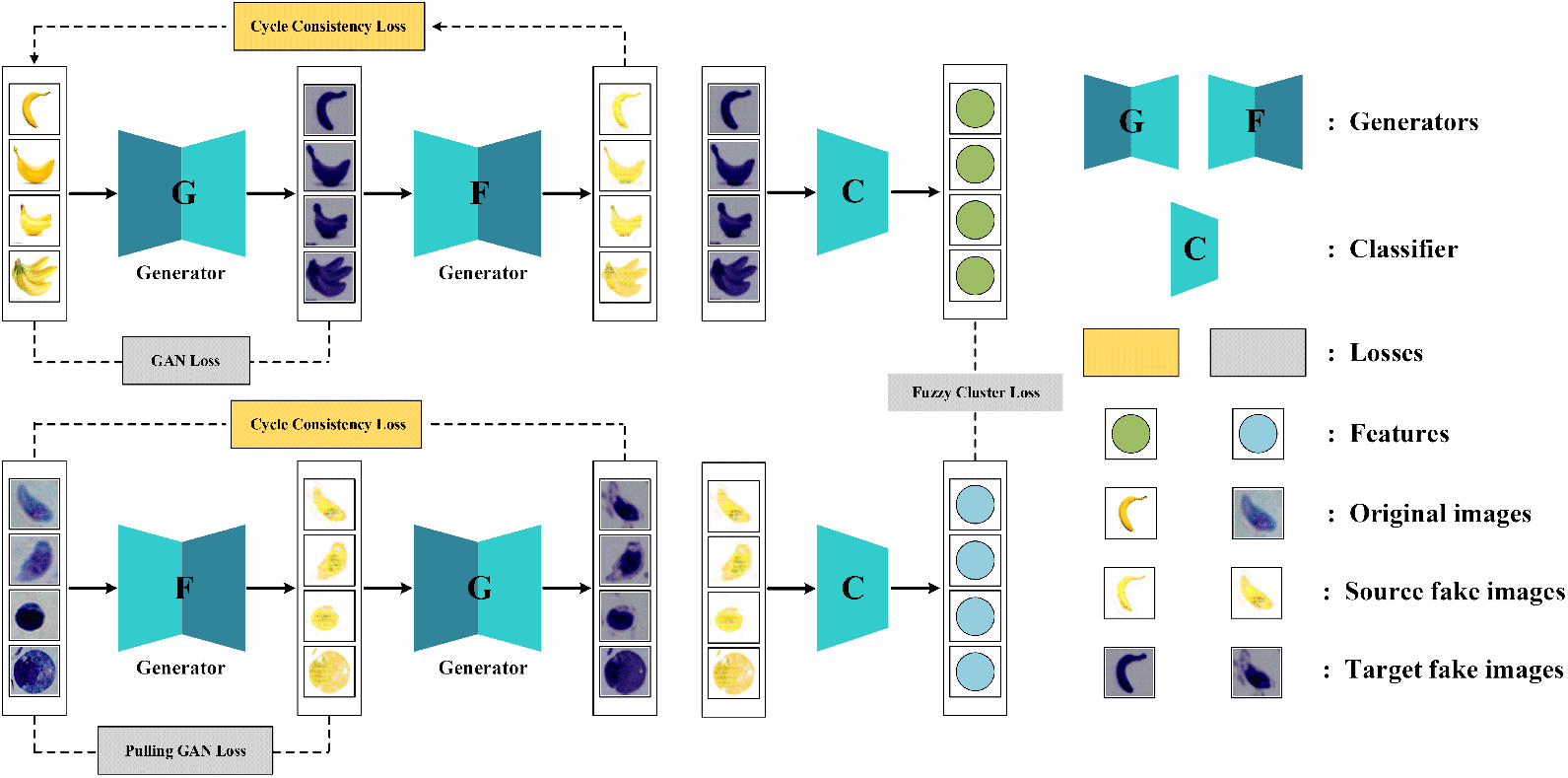
Schematic representation of FCGAN. Losses are shown in dashed rectangles. After training with source and target data alternatively, the generators and are used to pull *T.gondii* into source domain and keep host cells in the target domain according to their shape and texture information. Discriminators and are not shown in the architecture. Source fake images are the output of target images through generator and target fake images are the output of source image through generator. Features are extracted by a classifier for both images from source and target.

### 2.1 Preliminary Knowledge

To extract robust feature for microscopic images, we construct a deep neural network to compute the representation of each sample *x* by passing it to multiple layers of non-linear transformations. The key advantage of using such a network to map x is the nonlinear mapping function can be explicitly obtained. Assume there are *M* + 1 layers in the designed network and *p*^(*m*)^ units in the *m*-th layer, where *m* = 1, 2, …, *M*. The output of *x* at the *m*-th layer is computed as

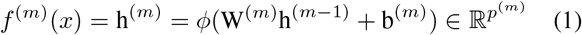

where 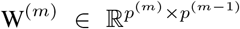 and 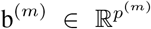 are the weight matrix and bias of the parameters in this layer, and ϕ is a nonlinear activation function which operates component-wisely, such as widely used *tanh* or *sigmoid* functions. The nonlinear mapping 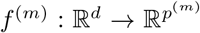 is a function parameterized by 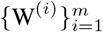 and 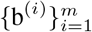. For the first layer, we assume h^(0)^ = x and *p*^(0)^ = *d*.

In this paper, we employ two typical convolutional networks as base framework in FCGAN, including Visual Geometry Group Network (VggNet) [Simonyan and Zisserman, 2014] and Cycle Generative Adversarial Network [Zhu *et al.*, 2017].

The VggNet architecture was introduced by Simonyan and Zisseman in their 2014 paper, Very Deep Convolutional Networks for Large Scale Image Recognition [Simonyan and Zisserman, 2014]. This network is characterized by its simplicity, using only 3 × 3 convolutional layers stacked on top of each other increasing depth. Reducing volume size is handled by max pooling. Two fully-connected layers, each with 4096 nodes are then followed by a soft max layer.

Cycle GAN is a Generative Adversarial Network (GAN) that uses two generators and two discriminators. We call one generator *G*, and have it convert images from the *X* domain to the *Y* domain. The other generator is called *F*, and converts images from *Y* to *X*. Each generator has a corresponding discriminator, which attempts to tell apart its synthesized images from real ones. Along with two components to Cycle GAN objective functions, an adversarial loss and a cycle consistency loss are essential to getting good results. Detail description can be seen in [Zhu *et al.*, 2017]

### 2.2 Micro-Macro Associated Object Pulling Strategy

A set of unlabeled microscopic images is classified as target domain 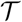 including *T.gondii* and host cells, another set of its associated macro object images is used as source domain, such as bananas. It is expected that source data shares the same cluster point with target *T.gondii* images, and host cell images belong to another cluster point. Then we calculate the degree of membership of each cluster point for each sample in 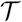. Furthermore, an image translator is used to pull the target *T.gondii* images closer to source banana images and the host cell images away from banana, which is achieved by replacing the domain label with the degrees of membership of each cluster point. Through the whole process, labeling of 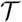 is not required and necessary. We integrate Cycle GAN to the image translator combined with a fuzzy C-means algorithm to enhance its selectivity, which discriminator labels are replaced by degrees of membership obtained in fuzzy C-means.

For a set of unannotated target domain 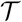 and its desired associated source domain 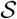, Cycle GAN learns two mappings without any supervision, 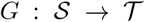 and 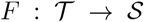 with two generators *G* and *F*, at the same time. To bypass the infeasibility of pixel-wise reconstruction with unpaired data, i.e. 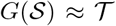 or 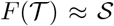, Cycle GAN introduces an effective cycle-consistency loss for 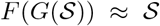 and 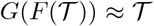. The idea is that the generated target domain data is able to return back to the exact data in the source domain where it generated from. To guarantee the fidelity of fake data 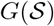 and 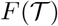, Cycle GAN uses two discriminators 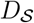 and 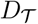 to distinguish real or synthetic data and thereby encourage generators to synthesize realistic data.

To solve the task of learning generators with unpaired images from two domains, and, we adopted the idea of the original cycle-consistency loss [Zhu *et al.*, 2017] for generators *G* and *F*, forcing the reconstructed synthetic sample 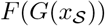 and 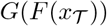 to be identical to their inputs 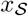 and 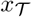.

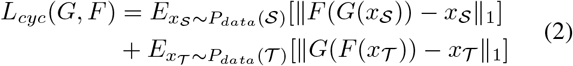

where 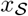 is the image from source domain 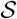 and 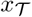 is from target domain 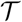. *L*_*cyc*_ uses the *L*_1_ loss, which shows better visual results than the *L*_2_ loss.

### 2.3 Challenges in M2AOP Strategy

Lacking supervision with a direct reconstruction error between 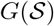 and 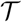 or 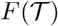 and 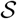 brings some uncertainties and difficulties towards the desired strengthened output of *T.gondii* and weakened output of host cells. The conventional Cycle GAN cannot evaluate the importance of each sample and it will transform all images in 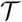 domain without any selectivity. That makes our task even more challenging.

Such problem cannot exploit the discriminative information from the associated object 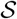 in and cannot select effective target samples from 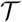 in an unsupervised manner. Our goal is to pull target *T.gondii* images closer to banana images in 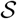 and make the other host cell images remain a large distance from source banana images. So, we design the FCGAN approach, consisting of source-target image translated consistency, fuzzy pulling cluster generator and discriminator, and its optimization.

### 2.4 Fyzzy Cycle Generative Adversarial Network

Aiming to obtain selectivity on Cycle GAN in *T.gondii* recognition without the help of microscopic annotations, we consider using cluster algorithm. Specifically, fuzzy clustering algorithm resolves this dilemma by introducing a degree of membership for each data point belonging to arbitrary number of clusters [Simon, 1996].

We embedded the degree of membership into Cycle GAN which denotes the degree of each sample belonging to *T.gondii* cluster point and host cell point. We assumed that *T.gondii* shares a same cluster point with banana images in, 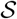 then the cluster closer to source banana point is the *T.gondii* cluster, and the other point represents host cell images. Then, we developed the generator loss especially for discriminators embedded with degrees of membership into adversarial loss. It could strengthen the generativity of embedded Cycle GAN for samples belonging to group, weaken the samples belonging to host cell group. As a result, the cluster points of *T.gondii* and host cell can be separated with a larger distance in the embedded Cycle GAN.

For this idea, we introduced the Fuzzy C-Means (FCM) algorithm [Bezdek, 1973]. To learn representative features, we designed a feature extractor *C* improved from Visual Geometry Group(VGG) network [Simonyan and Zisserman, 2014]. For images {*x*_1_, ⋯, *x*_*k*_, ⋯, *x*_*N*_} in target domain 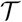 in parameter initialization (their translated fake source images are {*F*(*x*_1_), ⋯, *F*(*x*_*k*_), ⋯, *F*(*x*_*N*_)}), we can get their feature space after translation of {*C*(*F*(*x*_1_)), ⋯, *C*(*F*(*x*_*k*_)), ⋯, *C*(*F*(*x*_*N*_)} through the feature extractor *C*. Partition of these features into *P* clusters with minimization is defined as following objective function:

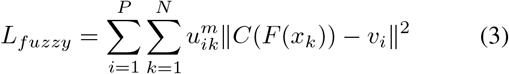

where, *x*_*k*_ is the *k*^*th*^ image in domain 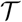, *v*_*i*_ is *d*-dimensional center of the *i*^*th*^ cluster, *u*_*ik*_ denotes the degree of membership of feature *C*(*F*(*x*_*k*_)) to cluster *v*_*i*_, *N* is the total number of features in each domain, ||*C*(*F*(*x*_*k*_)) − *v*_*i*_|| is any norm expressing the similarity between the measured feature *C*(*F*(*x*_*k*_)) and the cluster center *v*_*i*_. *m* denotes a weighting exponent parameter (*m* > 1) on each fuzzy membership value, and it determines the amount of fuzziness of the resulting classification according to:

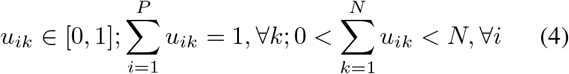

The first main step of this iterative algorithm is to update the membership function to determine in which cluster the feature belongs according to

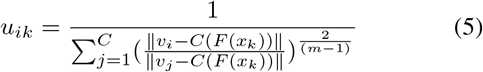

where *C* is a feature extractor.

The second main step concerns the updated centroids based on the new membership matrix according to

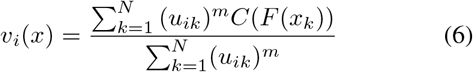

Finally, we compute the objective function related to Eq.3 and check the criteria termination (the objective function convergence).

The fuzzy membership matrix *U* consists the degree of membership belonging to each cluster for each sample feature. We embed *U*, which has two degrees belonging to *T.gondii* and host cell for each feature, denoted as 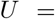 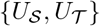, into our Cycle GAN, and design the fuzzy Cycle GAN loss for generators {*G, F*} and discriminators 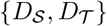. Their corresponding loss constraints are,

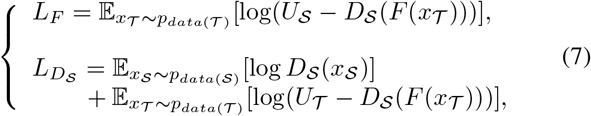

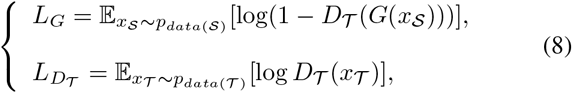

where 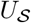 is the degrees of membership belonging to the shared cluster point of source banana images in 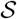 and *T.gondii* images in 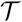, and 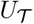 is the degrees of membership belonging to host cell cluster point in 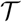. From these two constraints, 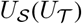 instead of label 1 in 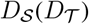 makes fake images from *T.gondii* images more similar to images 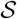 in than host cells images.

Algorithm 1 summarizes the main step of this sequential iterative algorithm with back-propagation and Lagrange Multiplier method.

### 2.5 Architecture and Training Detail

The architecture of our method is composed by original Cycle GAN and VGG layers. Cycle GAN originally designs generators with multiple residual blocks [He *et al.*, 2016], and discriminators with several convolutional layers combined with binary *L*2 loss [Zhu *et al.*, 2017], which setting is followed to maintain the generativity in Cycle GAN. Differently, in our discriminators, we make several critical modifications.

First, using VGG network [Simonyan and Zisserman, 2014] to learn feature representation is critical to maintain the discriminative information for microscopic images, as it has achieved much faster convergence and locally smooth results in malaria recognition [Mehanian *et al.*, 2017]. Second, we adopted the degrees of membership into the generative adversarial loss function to enhance the generativity of sample group belonging to the closed cluster between source domain, weaken the sample group belonging to the distant cluster between source domain for generators and discriminators. Such approach can separate the different group with a large distance.

We implemented our network on TensorFlow framework [Abadi *et al.*, 2016] with Tesla K40C and 128G memory in Ubuntu 16.04 system, and use the Adam solver [Kingma and Ba, 2015] for FCGAN network with a learning rate of 2*e* − 4 for generators, 2*e* − 6 for feature extractor, closely following the settings of Cycle GAN and VGG Network to train generators, discriminators and feature extractors. After jointly training for 4000 epochs, we apply early stop when the fuzzy loss no longer decreases for about 5 epochs (usually takes 4000 jointly training epochs to reach a desired point). In training, the number of training data in two domains can be different. Our parameters in Algorithm 1 are set as *m* = 2, *P* = 2 and *loop* = 100.

### 2.6 Image Collection and Evaluation

*T.gondii* microscopic images were captured under two brightfield light microscopes (Leica DM2700P and Olympus IX53), where preserved slices of parasites infection samples are mounted onto slides and stained with Giemsa. The first dataset is captured with 40X objectives (the magnification is 400X, T400) in Leica DM2700P microscope, obtaining 8,156 *T.gondii* images and 4,979 host cell images. The second dataset is captured with 100X oil immersion objectives (the magnification is 1000X, T1000) in Olympus IX53 microscope, which obtains 6,969 *T.gondii* images and 8,023 host cell images. In addition, we crawl 2,382 banana images, 2,053 crescent images and 1,860 star images on the Internet as different source data to compare macroscopic associated objects.

#### Algorithm 1 Fuzzy Cycle Generative Adversarial Network

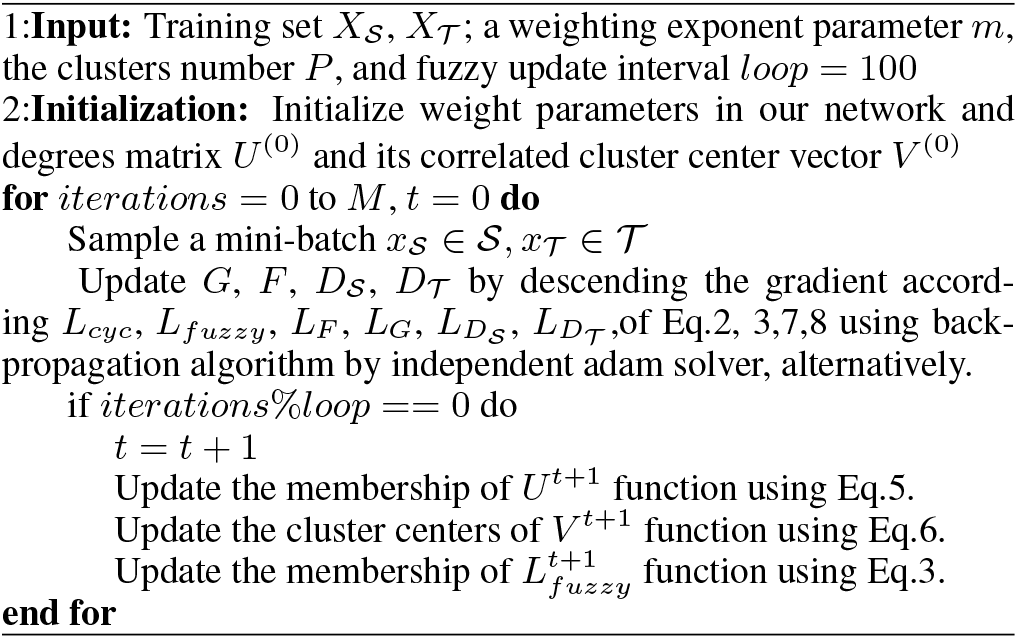

Notably, different magnification microscope slides typically display variations in color, background, and illumination between slides from different technicians, laboratories, clinics, and regions. Color variation can result from differences in staining pH, time and purity of dye, duration of the staining procedure, and the sensor settings (Figure 1). If uncorrected, these variations may degrade the model performance. To overcome negative effects of variations, we adopt white balancing techniques to compensate some of these variations. For all images, we use white balancing technique to pool the pixels from all fields of view and compute a global color balance affine transform for each image. Then we use a level-set based algorithm [Lankton and Tannenbaum, 2008] to crop *T.gondii* and host cells, and resize all images in 256 × 256 before feeding into the deep learning network.

To evaluate the overall recognition performance, we calculate the average recognition accuracy for our parasites datasets, and compute the F1-score, recall and precision for parasites and their host cells. The average values are summarized in Table 1. We also perform average area under the Receiver Operating Characteristic (ROC) curves for binary classification [Sirinukunwattana *et al.*, 2016], and the ROC curves are drawn in Figure 3. AUC measures the probability that given a pair of samples with different class label. If a sample from class, the classifier will be signed a high prediction score, compared to the samples from other classes. Here, the prediction score is calculated by its degree of membership.

**Table 1:** Results of FCGAN and baselines on T400 and T1000 Toxoplasma dataset.

| Dataset | T400 dataset |  |  |  | T1000 dataset |  |  |  |
| --- | --- | --- | --- | --- | --- | --- | --- | --- |
| Models | Accuracy | F1-score | Recall | Precision | Accuracy | F1-score | Recall | Precision |
| Supervised Learning Method |  |  |  |  |  |  |  |  |
| ResNet | 98.4% | 0.986 | 0.982 | 0.990 | 98.9% | 0.989 | 0.984 | 0.995 |
| VggNet | 98.7% | 0.989 | 0.996 | 0.982 | 99.1% | 0.991 | 0.990 | 0.991 |
| GoogLeNet | 97.2% | 0.974 | 0.950 | 0.998 | 99.6% | 0.997 | 0.999 | 0.993 |
| Unsupervised Learning Method |  |  |  |  |  |  |  |  |
| ResFCM | 84.3% | 0.851 | 0.817 | 0.889 | 80.0% | 0.801 | 0.803 | 0.800 |
| VggFCM | 77.4% | 0.760 | 0.650 | 0.916 | 87.6% | 0.863 | 0.785 | 0.960 |
| GoogLeFCM | 74.3% | 0.758 | 0.730 | 0.789 | 81.5% | 0.797 | 0.728 | 0.880 |
| CycleFCM | 86.9% | 0.889 | 0.955 | 0.832 | 86.2% | 0.862 | 0.861 | 0.863 |
| FuzzyFCM | 86.0% | 0.887 | 0.992 | 0.802 | 91.7% | 0.918 | 0.933 | 0.904 |
| GANFCM | 81.0% | 0.838 | 0.890 | 0.792 | 88.9% | 0.892 | 0.911 | 0.873 |
| **FCGAN+Star** | 74.8% | 0.767 | 0.749 | 0.786 | 87.0% | 0.867 | 0.845 | 0.889 |
| **FCGAN+Crescent** | 92.0% | 0.927 | 0.916 | 0.938 | 92.3% | 0.926 | 0.950 | 0.903 |
| **FCGAN+Banana** | **93.1%** | **0.939** | **0.960** | **0.919** | **94.0%** | **0.939** | **0.929** | **0.949** |

**Figure 3:**
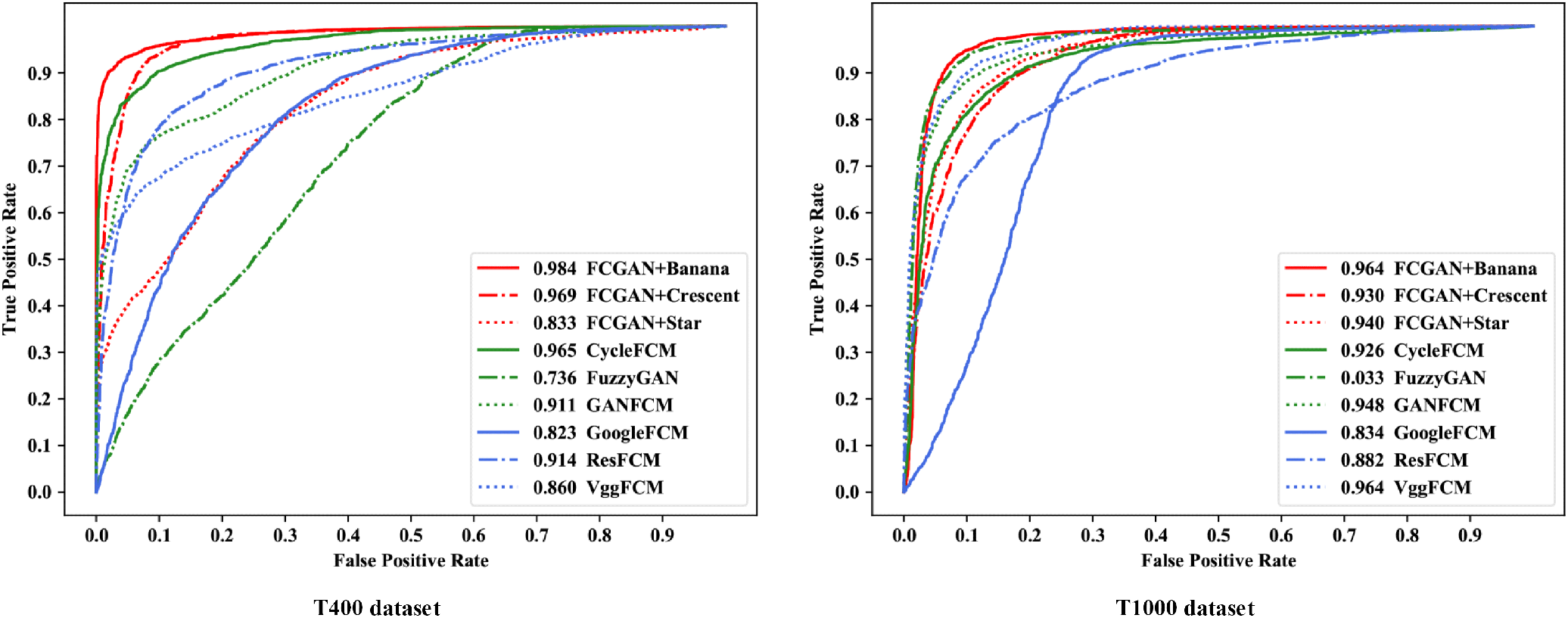
ROC curves of different methods for Toxoplasma recognition in T400 and T1000 datasets. The AUC value is labeled before the name of each method. FCGAN with banana source achieves the best performance compared to other unsupervised methods.

To validate the effectiveness of our FCGAN approach on *T.gondii* recognition task, we first compare FCGAN with three supervised deep Convolutional Neural Network (CNN) networks, including deep Residual Network (ResNet) [He *et al.*, 2016], Visual Geometry Group Network (VggNet) [Simonyan and Zisserman, 2014], Google Inception v4 network (GoogleNet) [Szegedy *et al.*, 2017]. We select half samples to train each supervised network and test on the left images for two datasets Besides, to discuss effects of components in our FCGAN, we implement some unsupervised experiments to evaluate effects of each component in FCGAN. VggFCM (ResFCM, GoogleFCM) is a combination of VGG (Res, Google) network with fuzzy C-means (FCM) cluster algorithm, and CycleFCM is combined the VGG+FCM and conventional Cycle Generative Adversarial Network (Cycle GAN), which can prove the efficiency of the embedded fuzzy component in Cycle GAN. Then, we also replace Cycle GAN with Generative Adversarial Network (GAN) [Radford *et al.*, 2015], and produce two baselines as fuzzyGAN with fuzzy domain labels, GanFCM with conventional GAN and FCM. Furthermore, we choose star and crescent as another source macroscopic objects besides banana, because star is not similar, but crescent is close to *T.gondii*.

To prove the main idea that the more similar of the macro object, the better performance will be achieved in the FC-GAN, we also perform extensive evaluations, including (a) cross domain image generation to test fuzzy component, (b) feature map visualization for convolutional layers, (c) t-SNE plots for feature extraction.

## 3 Result

### 3.1 Recognition performance of the model

As shown in the Table 1 of recognition performance in 400X and 1000X *T.gondii* datasets, our FCGAN with banana source data achieves the best performance in all unsupervised models with a small difference to supervised deep learning models (ResNet, VggNet and GoogleNet). By comparing with supervised methods, we select three widely used supervised CNN networks including VggNet, GoogleNet, and ResNet. As shown in Table 1, it can be seen that ResNet, GoogleNet and VggNet obtain 98.4%, 98.7% and 97.2% accuracy in T400 dataset. FCGAN with banana source achieves a competitive result of 93.1% accuracy, 0.939 F1-score, 0.960 recall, and 0.919 precision on T400 dataset, and 94.0% accuracy, 0.939 F1-score, 0.929 recall, and 0.949 precision on T1000 dataset. The values of accuracy, F1-score, precision and recall in our model are similar to those from ResNet, Vg-gNet and GoogleNet. All those results demonstrate our FC-GAN can achieve similar performance compared with supervised deep learning methods, while FCGAN do not require labeling of any microscopic images. It should be emphasized that *T.gondii* labeling under microscope is time-consuming, labor intensive and requires well trained professionals.

In comparison with unsupervised methods, we test five unsupervised baselines together with our FCGAN, including VggFCM, ResFCM, GoogleFCM, CycleFCM and GanFCM. The results are illustrated in Table 1, and Receiver Operating Characteristic (ROC) curves with AUC (area under the curve) value are drawn in Figure 3. As shown, it is easy to prove that our FCGAN achieves the best performance compared to other unsupervised methods, leaving the baselines at least 6.2% (93.1%-86.9%) accuracy, 0.05 (0.939-0.889) F1-score, 0.005 (0.960-0.955) recall, and 0.003(0.919-0.916) precision on T400 dataset.

To discuss the effectiveness of source object, we then compare the results from banana, crescent and star images as source. In morphology, star is less similar to *T.gondii* than banana and crescent, and its results and ROC curves have a distance between FCGAN+Banana and FCGAN+Crescent in Table 1 and Figure 3. The accuracy results are 93.1% (94.0%) of banana source, 92.0% (92.3%) of crescent source and 74.8% (87.0%) of star source on T400(T1000) dataset. It can be seen that FCGAN+Banana obtains a better performance than FCGAN+Crescent and FCGAN+Star in both dataset. Those results indicate the more similar of the macroscopic object, the better performance will be achieved in our FCGAN model.

Taken together, our Micro-Macro Associated Object Pulling strategy with banana source can achieve competitive results for *T.gondii*. recognition without any microscopic image labels.

### 3.2 Evaluation of the fuzzy component

In Figure 4, we illustrate a few samples of the translated images generated from FCGAN with different source objects and GAN-based baselines. FCGAN preserves most information in translated images including shape, nucleus and texture information, with the same background and color to their source objects. But for GAN-based methods, CycleFCM keeps most background information of images, and FuzzyGAN only generates target objects partially, missing some part of morphology information both in *T.gondii* and host cell. Similarly, GAN+FCM leaves out some morphology feature for host cell. Besides, compared with different source objects, FCGAN with banana source produces a clear outline without any background, compared to other source objects.

**Figure 4:**
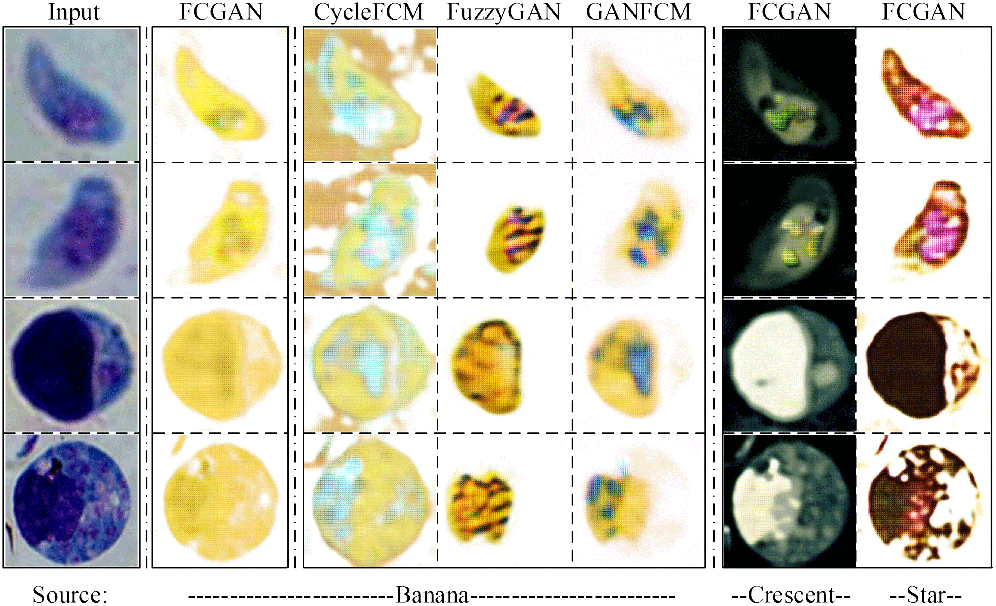
Image translation generated from different sources by FCGAN and other GAN-based methods. Compared to the GAN-based methods, the images generated from FCGAN contain more texture and shape details. Most notably, FCGAN with banana source has learned to reconstruct the nucleus of *T.gondii*.

Together, our FCGAN can create the texture and shape relationship between microscopic object and macroscopic object with selectivity of associated information. The comparison of different models can show that our FCGAN with banana source can achieve a better performance indicating FC-GAN can exploit associated information (for example, shape and texture information) to boost the cluster results.

### 3.3 Analysis of network layers

To analyze the network layers, we do feature map visualization for the first convolutional layer (Figure 5). Feature visualization is a key technique that helps identify and recognize patterns inside of neural networks. T-distributed Stochastic Neighbor Embedding (t-SNE) plots for the last layer are performed for all models (Figure 6). t-SNE is a non-linear dimensionality reduction approach that allows embedding high dimensional data in a lower-dimensional space.

**Figure 5:**
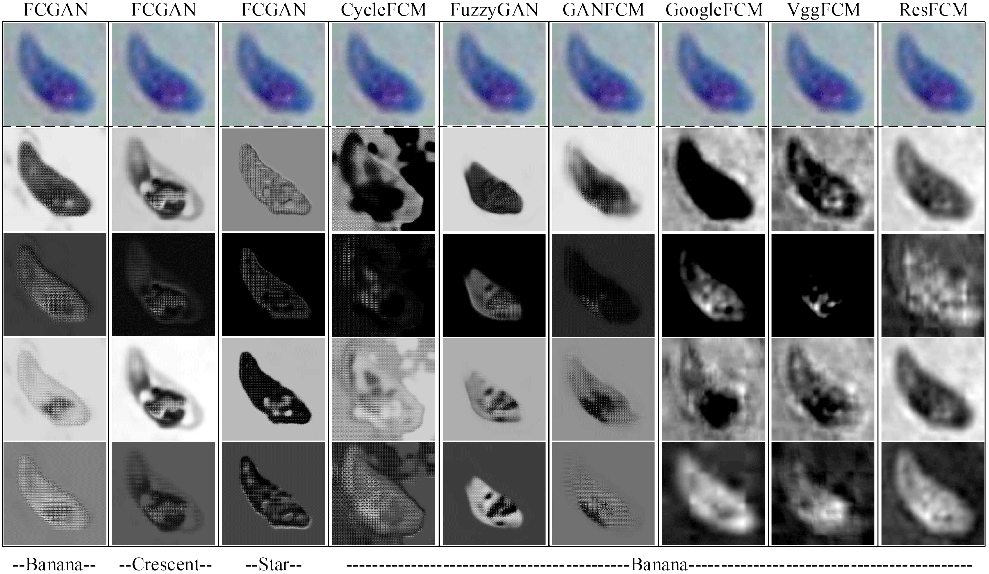
Visualization of the convolutional feature map learned by FCGAN and baselines. Object outline has been extracted and visualized for the first convolutional layer of FCGAN and other baselines.

**Figure 6:**
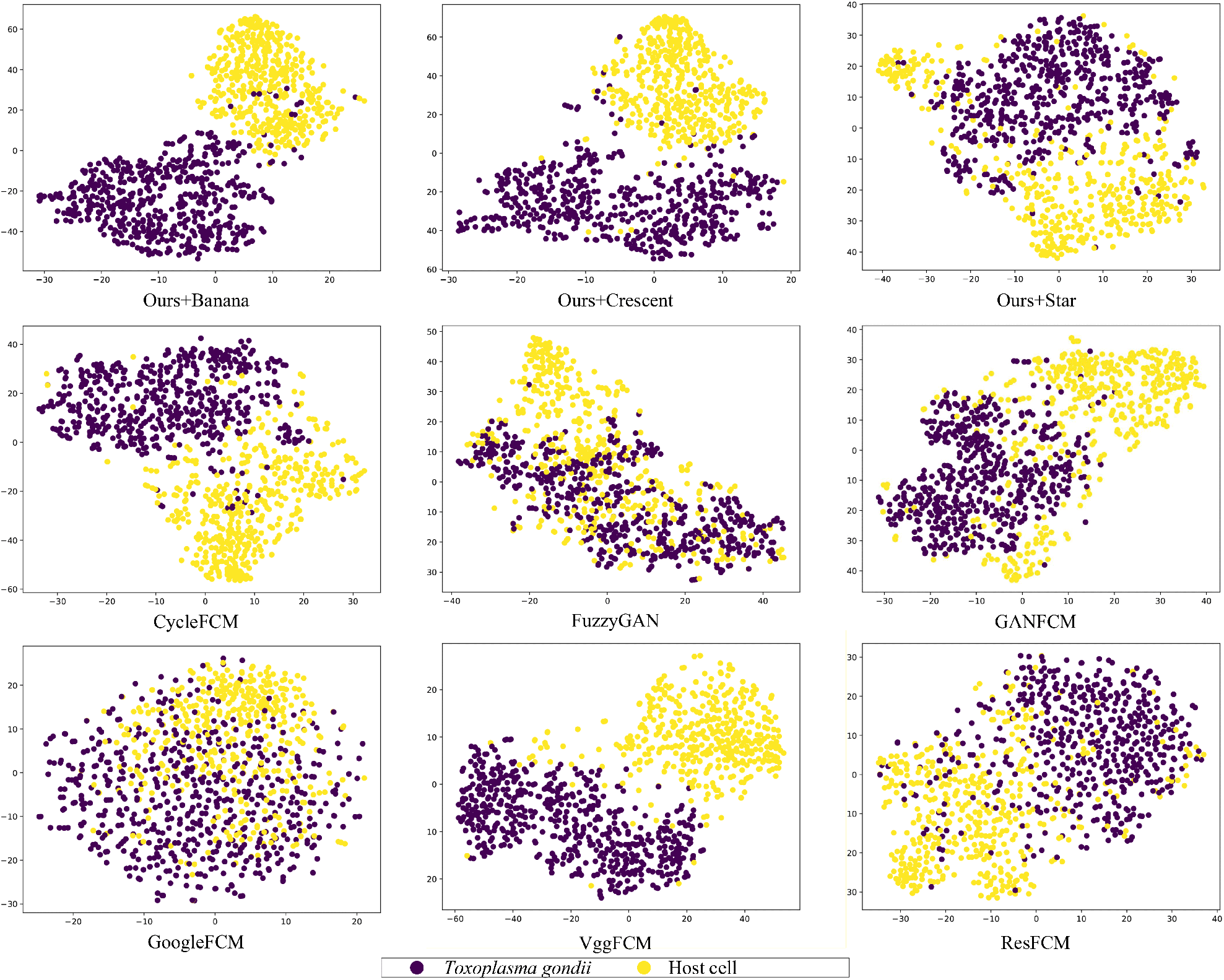
t-SNE plot of FCGAN and other baselines. t-SNE mappings provide a method to evaluate and refine clustering of parasite and cell images. Data points are colored according to cluster membership.

FCGAN can extract outline of target images with more clear morphology than other baselines. It demonstrates the FCGAN focus on their texture information and produce clear feature map for these information. FCGAN with banana source preserves the nucleus information in *T.gondii* image. Nucleus information of *T.gondii* is lost in crescent and star source. FCGAN achieves the best in t-SNE plot performance compared to other baselines. t-SNE plot shown the feature extracted by the last layer in FCGAN with banana source is the most discriminated. All these conclusions can prove that our FCGAN focuses on a clear outline of *T.gondii* in shallow convolutional layers and can extracts discriminative information in the network.

### 3.4 Application of the system for unstained *T.gondii* detection

To investigate if our AI system can be used in the diagnosis of other *T.gondii* microscopic images, we conduct the same transfer learning framework to analysis of unstained parasite images (Figure 1(c)). 500 unstained *T.gondii* and 500 unstained host cell images are collected under a bright field microscope to feed into our model. We achieve 90.3% accuracy, 0.910 f1-score, 0.986 recall and 0.846 precision on this new unstained dataset. This result shows our FCGAN works not only on large scale stained parasite dataset, but also on unstained one with a limited number of data.

Microscope staining is a key technique used to enable better visualization under the microscope. A variety of staining techniques can be used with light microscopy. Labeling and staining of parasite in microscope image has always been challenging. As a result, AI powered system for parasite recognition without any staining might open a new area for parasite detection that showed potential to overcome the drawbacks of staining techniques.

## 4 Discussion and conclusion

To our knowledge, this is the first study of *T.gondii* recognition task and transfer learning based Fuzzy Cycle Generative Adversarial Network approach. The model is inspired by the knowledge from parasitologists that *T.gondii* is in banana or crescent shaped form.

In this context, resemblances between macro images and micro images are exploited in order to train the dataset. We design a Micro-Macro associated object pulling strategy and propose the FCGAN method for *T.gondii* microscopic image recognition without any data annotation. This method uses the fuzzy clustering algorithm to help selectivity and optimize cycle-consistency and associated objects pulling loss.

Using two *T.gondii* microscopic image datasets with magnification of 400X and 1000X, we successfully demonstrate that our FCGAN model has stronger associated information selectivity and better pulling effect than the other deep learning approaches. We employ different source objects data with different similarity to *T.gondii*. The ranking of performance in different source objects is banana > crescent > star, which is consistent with their similarity to Toxoplasma in shape.

Our method achieves similar classification accuracy when separating *T.gondii* from regular host cells without using labeled Toxoplasma images to train the model, but instead using banana, crescent and star images. The proposed technique while obtaining worse results than established supervised techniques, outperforms unsupervised techniques by using an “analogy” macroscopic dataset instead of a real microscopic dataset of *T.gondii*.

However, the performance of our model depends highly on the macro object images looking like the objects in the microscopic images. Therefore, the performance of this model would likely be enhanced by testing on a more similar source image dataset. If this approach needs to be applied to another biomedical image domains, the macro object that best match the micro objects of interest is required.

In addition, the variation present in macro object photos is significantly influenced by the fact that they are 2D views of 3D objects whereas cells and parasites are flattened 3D objects with the sharpest focus image acquired. Therefore, multiple microscopic images taken at different focus distance need to be tested in the future.

Nevertheless, our method for Toxoplasma microscopic image analysis can potentially speed up the detection and pave the way for a rapid, low-cost diagnostics. Moreover, FCGAN can be applied to other biomedical image recognition tasks, which has complicated procedure in data collection and annotation. Our model can learn useful knowledge from macro related object without time consuming and labor intensive image annotation and labeling.

## Notes

#### Summary of Updates

We have update some experimental detail and supplement some experiments.

## References

Martín Abadi, Paul Barham, Jianmin Chen, Zhifeng Chen, Andy Davis, Jeffrey Dean, Matthieu Devin, Sanjay Ghemawat, Geoffrey Irving, Michael Isard, Manjunath Kudlur, Josh Levenberg, Rajat Monga, Sherry Moore, Derek Gordon Murray, Benoit Steiner, Paul A. Tucker, Vijay Vasudevan, Pete Warden, Martin Wicke, Yuan Yu, and Xiaoqiang Zheng. Tensorflow: A system for large-scale machine learning. In 12th USENIX Symposium on Operating Systems Design and Implementation, OSDI 2016, Savannah, GA, USA, November 2-4, 2016., pages 265–283, 2016.

Loddo Andrea, Di Ruberto Cecilia, and Kocher Michel. Recent advances of malaria parasites detection systems based on mathematical morphology. Sensors, 18(2):513–, 2018.

Gecer Baris, Aksoy Selim, Mercan Ezgi, Shapiro Linda G., Weaver Donald L., and Elmore Joann G. Detection and classification of cancer in whole slide breast histopathology images using deep convolutional networks. Pattern Recognition, pages S0031320318302577–.

James Christian Bezdek. Fuzzy mathematics in pattern classification. Ph. D. Dissertation, Applied Mathematics, Cornell University, 1973.

James C. Bezdek. Cluster validity with fuzzy sets. Journal of Cybernetics, 3(3):58–73, 1973.

Alison Burrells, Alessandra Taroda, Marieke Opsteegh, Gereon Schares, Julio Benavides, Cecile Dam-Deisz, Paul M. Bartley, Francesca Chianini, Isabella Villena, and Joke van der Giessen. Detection and dissemination of toxoplasma gondii in experimentally infected calves, a single test does not tell the whole story. Parasites & Vectors, 11(1):45, 2018.

Eric M. Christiansen, Samuel J. Yang, D. Michael Ando, Ashkan Javaherian, Gaia Skibinski, Scott Lipnick, Elliot Mount, Alison O’Neil, Kevan Shah, and Alicia K. Lee. In silico labeling: Predicting fluorescent labels in unlabeled images. Cell, 173(3):792–803, 2018.

Kaiming He, Xiangyu Zhang, Shaoqing Ren, and Jian Sun. Deep residual learning for image recognition. In 2016 IEEE Conference on Computer Vision and Pattern Recognition, CVPR 2016, Las Vegas, NV, USA, June 27-30, 2016, pages 770–778, 2016.

Asis Khan and Michael E. Grigg. Toxoplasma gondii: Laboratory maintenance and growth. Current Protocols in Microbiology, 44(1):20C.1.1, 2017.

Diederik P. Kingma and Jimmy Ba. Adam: A method for stochastic optimization. In 3rd International Conference on Learning Representations, ICLR 2015, San Diego, CA, USA, May 7-9, 2015, Conference Track Proceedings, 2015.

Shawn Lankton and Allen R. Tannenbaum. Localizing region-based active contours. IEEE Trans. Image Processing, 17(11):2029–2039, 2008.

Courosh Mehanian, Mayoore Jaiswal, Charles Delahunt, Clay Thompson, and David Bell. Computer-automated malaria diagnosis and quantitation using convolutional neural networks. In IEEE International Conference on Computer Vision Workshop, 2017.

Alec Radford, Luke Metz, and Soumith Chintala. Unsupervised representation learning with deep convolutional generative adversarial networks. Computer Science, 2015.

D Simon. Fuzzy sets and fuzzy logic: Theory and applications. Control Engineering Practice, 4(9):1332–1333, 1996.

Karen Simonyan and Andrew Zisserman. Very deep convolutional networks for large-scale image recognition. Computer Science, 2014.

Korsuk Sirinukunwattana, Shan Raza, Yee-Wah Tsang, David Snead, Ian Cree, and Nasir Rajpoot. Locality sensitive deep learning for detection and classification of nuclei in routine colon cancer histology images. IEEE Transactions on Medical Imaging, 35(5):1196–1206, 2016.

Rajaraman Sivaramakrishnan, Antani Sameer K., Poostchi Mahdieh, Silamut Kamolrat, Hossain Md. A., Maude Richard J., Jaeger Stefan, and Thoma George R. Pre-trained convolutional neural networks as feature extractors toward improved malaria parasite detection in thin blood smear images. Peerj, 6(1):e4568-, 2018.

Christian Szegedy, Sergey Ioffe, Vincent Vanhoucke, and Alexander A. Alemi. Inception-v4, inception-resnet and the impact of residual connections on learning. In Proceedings of the Thirty-First AAAI Conference on Artificial Intelligence, February 4-9, 2017, San Francisco, California, USA., pages 4278–4284, 2017.

Fuyong Xing, Yuanpu Xie, Hai Su, Fujun Liu, and Lin Yang. Deep learning in microscopy image analysis: A survey. IEEE Transactions on Neural Networks & Learning Systems, PP(99):1–19, 2017.

Jun Yan Zhu, Taesung Park, Phillip Isola, and Alexei A. Efros. Unpaired image-to-image translation using cycle-consistent adversarial networks. In IEEE International Conference on Computer Vision, 2017.

